# G-Protein coupled Purinergic P2Y12 receptor interacts and internalizes Tau^RD^-mediated by membrane-associated actin cytoskeleton remodelling in microglia

**DOI:** 10.1101/2021.05.24.445460

**Authors:** Hariharakrishnan Chidambaram, Rashmi Das, Subashchandrabose Chinnathambi

## Abstract

In Alzheimer’s disease, the microtubule-associated protein, Tau misfolds to form aggregates and filaments in the intra- and extracellular region of neuronal cells. Microglial cells are the resident brain macrophage cells that are involved in constant surveillance and are activated by the extracellular deposits. Purinergic receptors are involved in chemotactic migration of microglial cells towards the site of inflammation. In our recent study, we found that microglial P2Y12 receptor has been involved in phagocytosis of full-length Tau species such as monomers, oligomers and aggregates by actin-driven chemotaxis. In this study, we have showed the interaction of repeat-domain of Tau (Tau^RD^) with microglial P2Y12 receptor and analysed the corresponding residues for interaction by various *in-silico* approaches. In cellular studies, Tau^RD^ was found to interact with microglial P2Y12R and induces its cellular expression as confirmed by coimmunoprecipitation and western blot analysis respectively. Similarly, immunofluorescence microscopic studies emphasized that Tau^RD^ were phagocytosed by microglial P2Y12R *via* the membrane-associated actin remodelling as filopodia extension. Furthermore, the P2Yl2R-mediated Tau^RD^ internalization has activated the microglia with an increase in the Iba1 level and Tau^RD^ become accumulated at peri-nuclear region as localized with Iba1. Altogether, microglial P2Y12R interacts with Tau^RD^ and mediates directed migration and activation for its internalization.

## INTRODUCTION

Alzheimer’s disease is characterized by aggregates of Tau and β-amyloid proteins in the intra- and extracellular regions of neuronal cells respectively (1). Amyloid-β, the cleavage product of amyloid precursor protein (APP) accumulates in the extracellular region as senile plaques (2,3). Tau is a microtubule-associated protein that aggregates as oligomers and filaments in the AD brain (2,4). Tau is a soluble, cytoplasmic protein, extensively expressed in neuronal cells for microtubule-binding and stability (4,5). Tau is expressed in six different isoforms with the longest isoform comprising of 441 amino acids with two N-terminal inserts, N1 and N2, proline-rich domains and a repeat domain with four repeats, R1, R2, R3 and R4 (6,7). The repeat domain forms the stable β-sheet structure and rapidly aggregates to form stable filaments (8).

Microglia are the resident immune cells that maintains constant surveillance in the central nervous system. Microglia are activated by molecules from damages neurons, cell debris, aggregated proteins, etc., (9). Several GPCRs and other membrane receptors are involved in the activation of microglia, which ultimately leads to pro- and anti-inflammatory responses (10–12). G-protein coupled receptors (GPCRs) such as formyl peptide receptor 2 (FPR2) and chemokine-like receptor 1 (CMKLR1) also act as receptors for microglial activation and phagocytosis (13,14). The accumulated amyloid-β peptides and Tau species activates microglia through a wide variety of membrane receptors. Amyloid-β receptors in microglia includes scavenger receptors such as SCARA-1, SCARB-1, MACRO & RAGE, toll-like receptors such as TLR-2 and TLR-4 and Triggering receptor expressed in myeloid cells (TREM-2) (13,14). Though there are not many evidences for Tau receptors, microglial internalization of Tau species such as monomers and oligomers have been well studied(15–17). In addition to microglia, astrocytes are also reported to internalize monomeric Tau to form intracellular aggregates (18). Recent report emphasized that extracellular Tau are internalized by microglia via interacting with heparan sulphate proteoglycan (HSPGs) while, astrocytes phagocyted Tau *via* HSPGs independent mechanism (18). The mechanism of internalization for monomeric vs aggregated Tau follows two distinct but inter-connected pathways. Aggregated Tau usually internalized *via* dynamin-dependant pathway related to endocytic mechanism. While, the internalization of monomeric Tau follows through slow actin-dependant micropinocytosis as well as endocytic pathway (19). In addition, Tau interaction is reported with a microglial GPCR, chemokine CX3C receptor-1 (CX3CR1) for its activation and internalization (15,20).

Purinergic receptors are membrane receptors that are expressed in microglia for its chemotactic migration towards purinergic molecules such as ADP, ATP, UDP, AMP, etc., P2Y12R is a purinergic GPCR that plays a vital role in platelet functions and haemostasis. In CNS, P2Y12R is abundantly expressed in microglial cells that are involved in constant surveillance and activation (21). It has been recently reported that microglia form P2Y12R-mediated somatic synapses with neurons in order to survey neuronal health (22). In physiological condition, microglia maintain the ‘ramified’ state with long processes, no net displacement to survey constantly the microenvironment. While, upon pathogenic attack, neuronal injury, protein aggregation activates microglia to retain the extensions and transformed into ‘ameboid’ state (23). Activated microglia follow the chemical gradient to migrate at the site of neuronal injury. Previously, our group has reported that microglia can phagocytose extracellular Tau oligomers and monomers via membrane-associated actin remodelling (24). Moreover, extracellular Tau exposure activate the microglia with increased Iba1 level and its colocalization with modified actin network (24). P2Y12 receptor involves p38 mitogen-activated protein kinase (MAPK) and ROCK2 kinase pathway for inflammatory and neuropathic pain (21,25,26). Moreover, P2Y12R mediates an multi-faced cellular signalling cascades which involves chemotactic migration, actin remodelling, inflammasomes, microglial activation and also in the maintenance of neuronal health (27). In our previous work, microglial P2Y12 receptor is found to interact with extracellular fulllength Tau species such as monomers and oligomers. The interaction of different fulllength Tau species with P2Y12 receptor promotes actin remodelling via the formation of lamellipodia, filopodia and the microtubule-organizing center (MTOC) localization for directed-mediated microglial chemotaxis migration in ATP-dependant manner (28).

In our present study, we report the domain responsible for direct P2Y12 receptor interaction and activation. The repeat-domain of Tau is extensively studied for its role in P2Y12R interaction, receptor-associated Tau^RD^ internalization, accumulation and microglial activation *via* the involvement of Iba1 expression. Similarly, the presence of P2Y12R in the formation of filopodia and podosome-like structures as components of membrane-associated actin remodelling were also emphasized upon Tau^RD^ exposure during microglial migration.

## RESULTS

### Molecular modelling and docking of repeat-domain of Tau with the purinergic receptor, P2Y12R

Since, Tau is unstable, non-structural and a highly soluble protein, Tau^RD^ domain was used here for the interaction studies (Fig. 1A, B). Repeat-Tau model was adopted from Sonawane et al. 2019 that has a stable secondary structure for hexapeptide regions ^275^VQIINK^280^ and ^306^VQIVYK^311^and plays a key role in aggregation and filament formation (Fig. 1B). The model is built for Tau residues 244-373 followed by structural validation and energy minimization (Fig. 1C). The antagonist-bound structure (inactive form) of P2Y12 receptor (PDB ID. 4NTJ), was used as template for the modelling and the best model has been used for further analysis. The predocking refinement is performed that includes adding hydrogens, building the missing residues, loops and the GPCR refinement in order to attain the stable structural conformation. The final refined model had an RMSD value of 0.4 Å and Ramachandran favoured residues of 98.2%. The extracellular, cytosolic and transmembrane domains were segregated based on UniProt database (Entry no. Q9H244) (Fig. 1D). Tau^RD^ model was docked to the extracellular domain of P2Y12 receptor (highlighted in red, fig. 1D) and the best model with a center score of −1359.2 and lowest energy score of −1857.4 is carried forward for the molecular dynamics simulation (Fig. 1E). Similarly, P2Y12R model for active form (4PXZ) and inactive form bound to antagonist (4NTJ with antithrombic drug) were also built and Tau^RD^ was docked to the respective extracellular domain of P2Y12 (data not shown).

**Figure 1.**
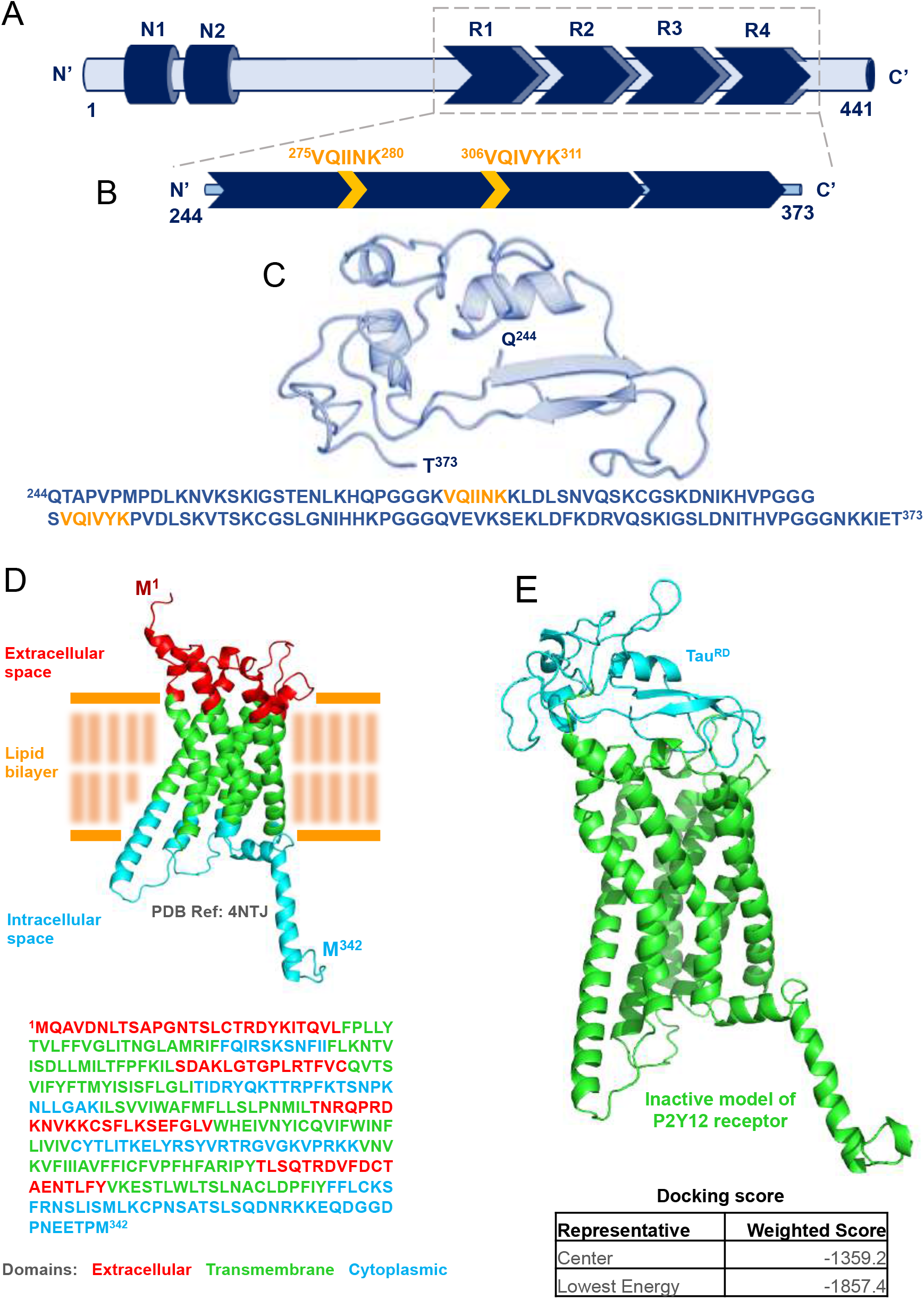
Molecular modelling and docking of repeat domain of Tau with the Purinergic receptor, P2Y12. **Structural characterization of Tau^RD^ and P2Y12 receptor. A.** Bar graph representing the full-length Tau (hTau40wt) structure of 441 amino acids with two N-terminal inserts (N1 and N2), and a repeat-domain comprising of four repeats (R1, R2, R3 and R4). **B.** Tau^RD^ comprising of two hexapeptide regions ^275^VQIINK^280^ and ^306^VQIVYK^311^ which are key residues in Tau aggregation. **C**. Tau model for the repeat domain, built using modeller with NMR structure, 2MZ7 as a template (Structure adopted from Sonawane *et al* 2019). The corresponding amino acid sequence of Tau^RD^ used for modelling studies. **D.** Diagrammatic representation of inactive form of P2Y12 receptor (PDB ID: 4NTJ) with phospholipid bilayer. Red denotes residues of extracellular domain; green denotes transmembrane residues and cyan denotes residues of cytoplasmic domain. The corresponding amino acid sequence of human P2Y12 receptor. **E.** Molecular docking model of P2Y12 receptor and Tau^RD^, with Tau^RD^ docked to extracellular domain of P2Y12 receptor (Green-P2Y12 receptor, cyan-Tau^RD^).

### Interaction studies of P2Y12R-Tau^RD^ by molecular dynamics simulation

Tau^RD^ complex with different models of P2Y12 receptor (active, inactive, and inactive with antagonist) were individually subjected to 200 nanoseconds simulation and analysed for the Lennard Jones potential and Coulombic energy between the interacting proteins (SI fig. 1). 4NTJ model of P2Y12R was chosen for further MD-simulation analysis. The P2Y12R - Tau^RD^ complex was subjected to a molecular dynamics simulation of 500 nanoseconds in order to interpret the stability, type of interaction and propose the corresponding amino acid residues involved in this interaction. Snapshot of the interacting complex at every 100 nanoseconds interval is shown in SI fig. 2. The stability of this interaction is predicted using root mean square deviation (RMSD) analysis with timescale (ns) and RMSD values (nm) along X- and Y-axis respectively. The complex (black) attained stability over the initial 100 ns and retained at 1 nm throughout the 500 ns simulation with a minimum deviation of 0.22 nm (Fig. 2B). Similarly, individual RMSD values of Tau^RD^ (green) and P2Y12R (red) were plotted over the 500 ns timescale (Fig. 2B). Since Tau is highly unstable, we checked for the fluctuating residues during the course of simulation through root mean square fluctuation (RMSF) analysis. The lysine residues such as K274, K290, K294, K298, K369 and K370 are showing peak fluctuations, whereas the hexapeptide regions are showing least fluctuations throughout the simulation (Fig. 2C). The RMSF graph shows the Tau^RD^ residual fluctuation at different time intervals, i.e., 0-100 ns (black), 200-300 ns (green) and 400-500 ns (magenta). The residual fluctuation reduced and attained stability during the latter part of the simulation (400-500 ns). We next plotted the energy graph for analysing, electrostatic, Vander Waal’s and total interaction energy between P2Y12R and Tau^RD^ complex. The total binding energy attains more negative during the latter part of simulation (350-500 ns) and attained around - 1250 KJ/mol during the course of simulation, which indicates a favourable interaction (Fig. 2D). We have calculated the number of hydrogen bonds formed in order to check the hydrogen bond interaction between the complex. The number of hydrogen bonds (black) reached a maximum of 27 during the initial timescale and attained a stable count around 10-15 during the simulation of 500 ns (Fig. 2E). Tau is already reported to interact with cellular membrane components. In this study, the Tau^RD^ also interacts with palmitoyl oleoyl phosphatidyl choline (POPC) membrane and the corresponding hydrogen bond analysis is mentioned in Fig. 2F with around 20-25 hydrogen bonds during the time of 350-500 ns of the simulation.

**Figure 2.**
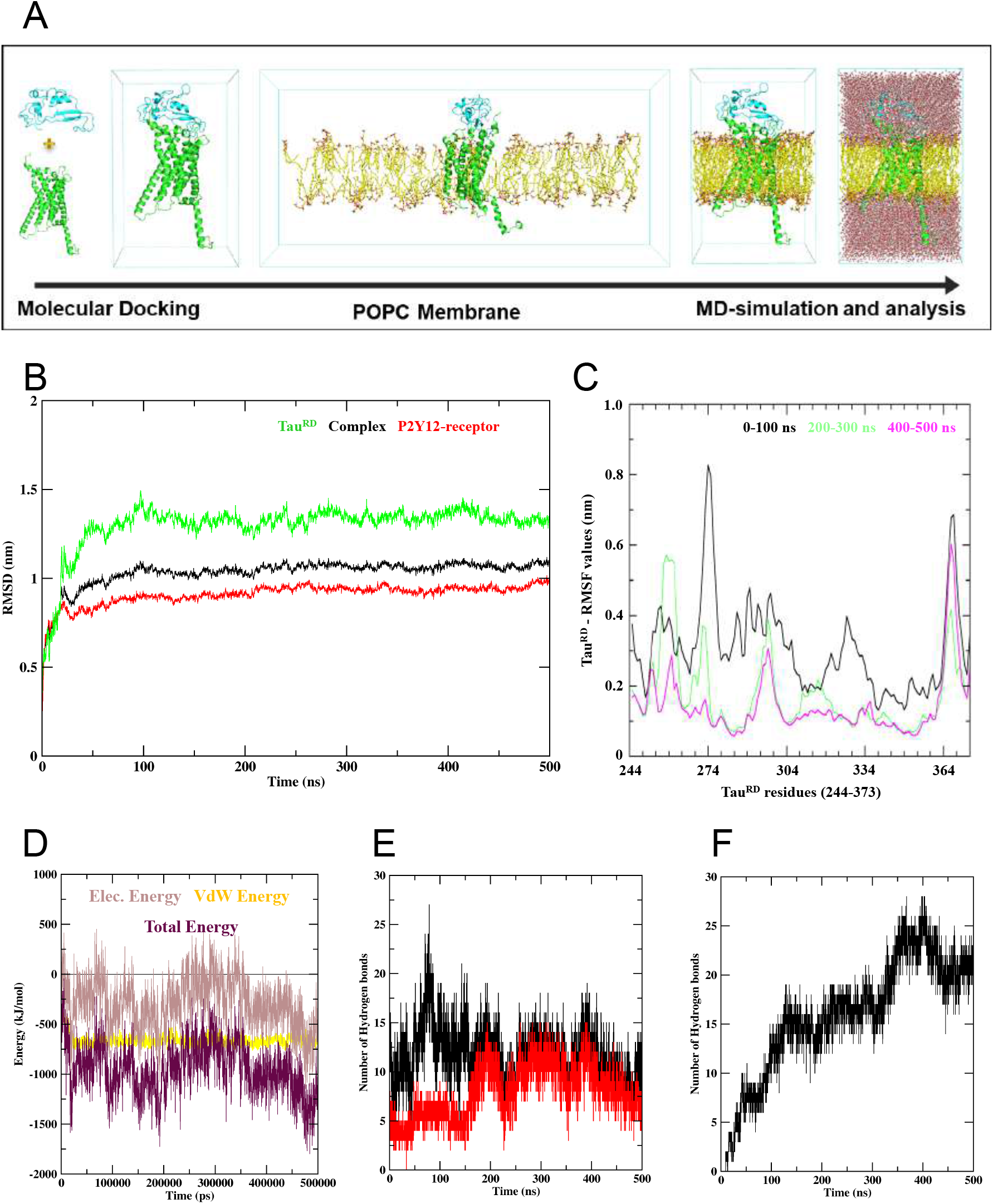
Molecular-Dynamics simulation of P2Y12R in complex with Tau^RD^. **A.** Outline of molecular docking and MD-simulation studies performed for P2Y12R-Tau^RD^ interaction studies. Tau docked to the extracellular domain of P2Y12R, surrounded by phospholipid bilayer and solvated with water, is subjected to molecular dynamics simulations. The overall simulation was performed for a time-scale of 500 nanoseconds. **B.** The root-mean square deviation (RMSD) graph showing time (ns) and RMSD values (nm) on the X- and Y-axis respectively. Green denotes RMSD values of Tau^RD^, red denotes P2Y12 receptor and black denotes P2Y12R-Tau^RD^ complex. **C.** The root-mean square fluctuation (RMSF) graph of Tau^RD^ showing RMSF values (nm) and amino acid residues on the Y- and X-axis respectively. Black denotes RMSF values of Tau^RD^ over 0-100 ns, green denotes 200-300 ns and magenta denotes 400-500 ns. **D.** Interaction energy graph (Extracellular domain of P2Y12R vs Tau^RD^) plotted with energy values (KJ/mol) and time-scale (ps) on Y- and X-axis respectively. Brown represents electrostatic energy, yellow represents Vander Waal’s energy and maroon represents the total energy between P2Y12R extracellular domain and Tau^RD^. **E.** Hydrogen bond analysis with number of hydrogen bonds plotted over time-scale (ns) on Y- and X-axis respectively. Black denotes total hydrogen bonds formed between P2Y12 receptor and Tau^RD^, whereas red denotes hydrogen bond formed between specific residues of Tau and P2Y12 receptor (mentioned in h-bond analysis of figure 4) that were stable throughout simulation. F. Hydrogen bond formed between Tau^RD^ and POPC membrane is plotted with number of hydrogen bonds and time-scale (ns) on Y- and X-axis respectively.

### Residual interaction analysis of P2Y12R-Tau^RD^ complex

For this part of the study, the trajectory of 350-500 ns simulation was considered and the residues that has a stable interaction (hydrogen bonding and hydrophobic interaction) has been plotted for residual interaction analysis (Fig. 3A). The charged amino acids of Tau are playing a key role in P2Y12R interaction. The amino acid residues such as D252, E269, K331, K343 and H330 of Tau^RD^ are contributing to the hydrogen bonding (plotted in Fig. 3B) with the extracellular residues of P2Y12R, K174, E181, D20, D269 and C17 respectively. Hence, the cysteine residues of P2Y12R, C17 and C270 could possibly play a critical role in P2Y12R-Tau^RD^ interaction. The contribution from these specific residues to the overall hydrogen bonding is plotted as red in Fig. 2E. The valine and proline residues of Tau such as V248, V339, V350, P247, P249 and P251 are majorly contributing to the hydrophobic interaction between the complex (Fig. 3A). Similarly, the extracellular residues of P2Y12R such as V172, K173, T260, F177, L87 and L184 are majorly contributing to the hydrophobic interactions with Tau^RD^.

**Figure 3.**
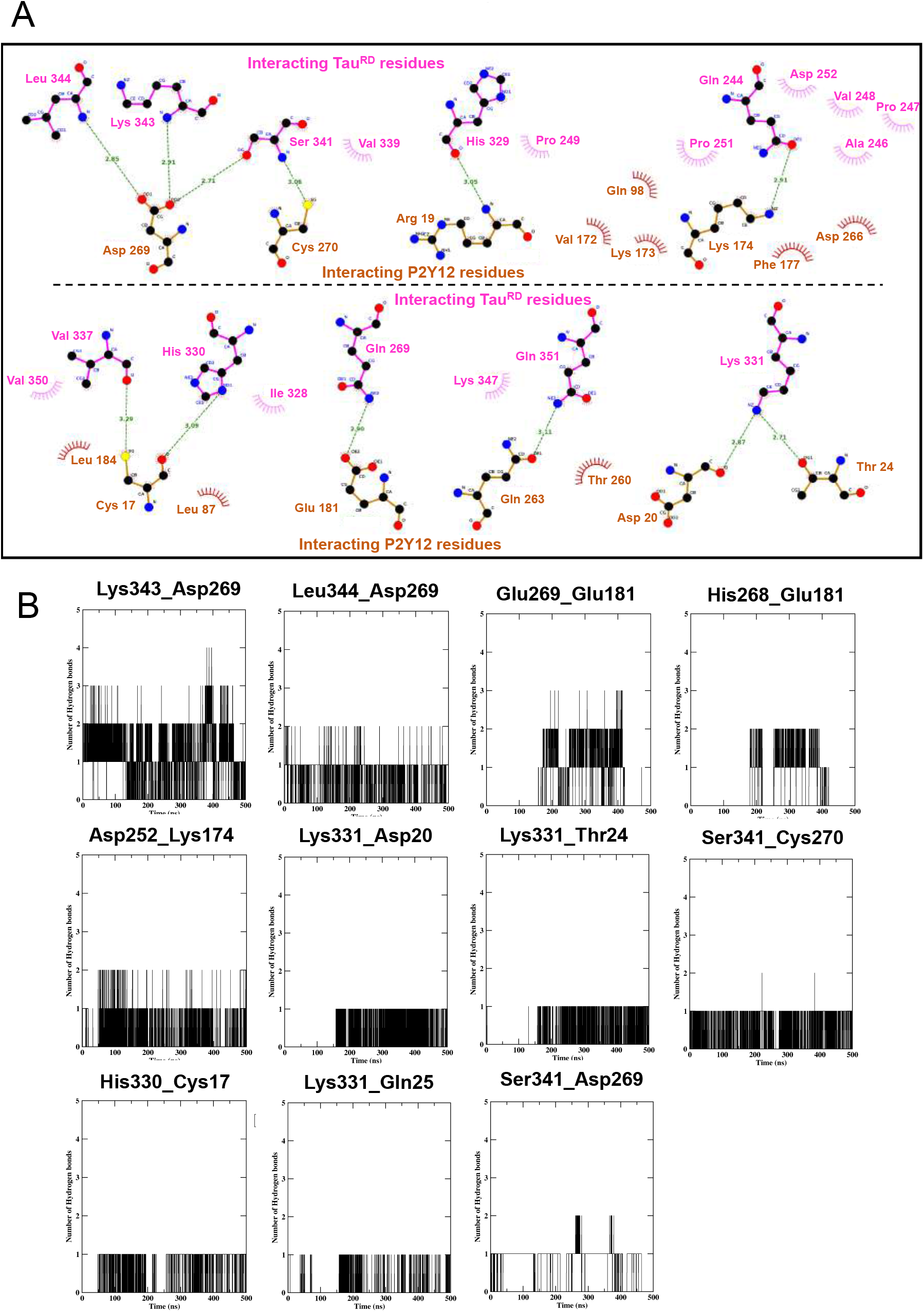
Residue-specific interaction analysis of P2Y12R-Tau^RD^ complex. **A.** 2D interaction analysis of P2Y12-Tau^RD^ that includes hydrogen bonding and hydrophobic residues (Data collected from the last 350-500 ns of simulation). **B.** Residual hydrogen bond analysis of Tau^RD^ with extracellular P2Y12 receptor that are identified from the 2D interaction graph with number of hydrogen bonds formed and time (ns) on Y- and X-axis respectively (Amino acid residue names on left and right side of the individual graph title corresponds to Tau^RD^ and P2Y12 receptor respectively).

We have also studied the surface electrostatic potential of the P2Y12R-Tau^RD^ complex obtained after 500 nanoseconds simulation (Fig 4). The protein colour is based on the electrostatic potential of protein surface where blue and red denotes the positive and negative energies (KJ/mol) respectively. The side view of the Tau^RD^ protein and a 90° rotation (bottom view) shows its surface potential on its interacting region (Fig. 4A), which is predominantly positively charged (blue). The surface potential of the interacting P2Y12R-Tau^RD^ complex and the side and top view is also visualized (Fig. 4B). In order to visualize the extracellular binding site of P2Y12 receptor for Tau^RD^ binding, the top view of the P2Y12R-Tau^RD^ complex with a cartoon representation for Tau^RD^ (green) is generated (Fig. 4C). This clearly suggests that Tau^RD^ interacts with the negatively charged N-terminal region of the extracellular P2Y12 receptor. The side view of the P2Y12 receptor and a 90°rotation (top view) shows the Tau^RD^-interacting surface, along the negatively charged N-terminal region (red) of the extracellular P2Y12 receptor which suggests a favourable interaction for the P2Y12R-Tau^RD^ complex. (Fig. 4D).

**Figure 4.**
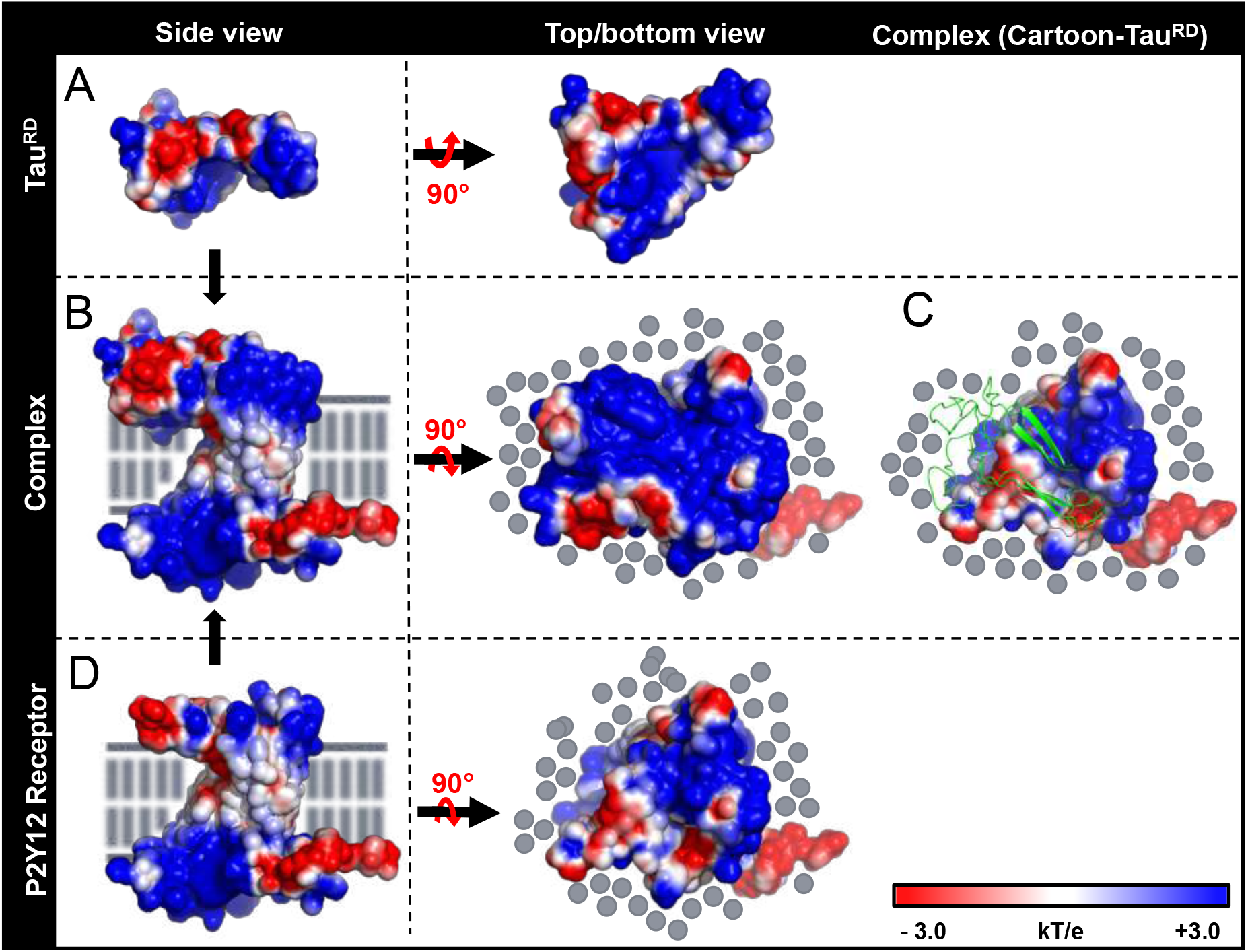
Surface electrostatic potential analysis of Tau^RD^ (A), P2Y12R-Tau^RD^ complex (B) and P2Y12R (C). Studies were performed from the P2Y12R-Tau^RD^ complex structure obtained after 500 nanoseconds simulation. The surface of the protein is coloured based on electrostatic potential where blue and red denotes the positive and negative energies (KJ/mol) respectively. Grey lines/ dots represent the phospholipid membrane surrounding P2Y12R. **A.** Side view of the Tau^RD^ and a 90° rotation (bottom view) shows the P2Y12R-interacting surface of Tau^RD^ that is predominantly positive. **B.** Side view of the P2Y12R-Tau^RD^ complex and a 90° rotation which shows the top view of the complex. **C.** Top view of P2Y12R-Tau^RD^ complex with cartoon representation for Tau^RD^ (green) that shows its binding region on the extracellular surface of P2Y12 receptor. **D.** Side view of the P2Y12 receptor and a 90°rotation (top view) shows the Tau^RD^ interacting surface, towards the negatively charged N-terminal region (red) of P2Y12 receptor.

### Activated microglia phagocytose Tau^RD^ and colocalize with Iba1

Microglia is one of most important brain-resident immune cells, which become activated by encountering extracellular protein deposits. Iba1 is a calcium receptor-binding adaptor protein which is involved in microglial activation and migration. In order to check microglial activation, N9 cells were exposed with Tau^RD^ for 24 hours. Upon encountering Tau^RD^, microglia have increased the level of Iba1 as observed by western blot and quantification (Fig. 5A, B). Moreover, when microglia were treated with Alexa 647-labelled Tau^RD^ for 24 hours, N9 cells have phagocytosed Tau^RD^ which was colocalized with Iba1, as observed by immunofluorescence study (Fig. 5C). The Pearson’s coefficient (R) of fluorescence of Iba1 and phagocytosed Tau^RD^ was quantified in microglia which ranges from 0.76-0.9 units which signify positive correlation in Tau^RD^-Iba1 colocalization (n=12) (Fig. 5D). Microglial activation was also studied by colocalization of Iba1 with migratory actin structures. Upon exposure of Tau^RD^, microglia remodelled membrane-associated actin network (F-actin) which was colocalized with Iba1, signifying microglial migration upon activation (Fig. 5E; SI fig. 3A and B).

**Figure 5.**
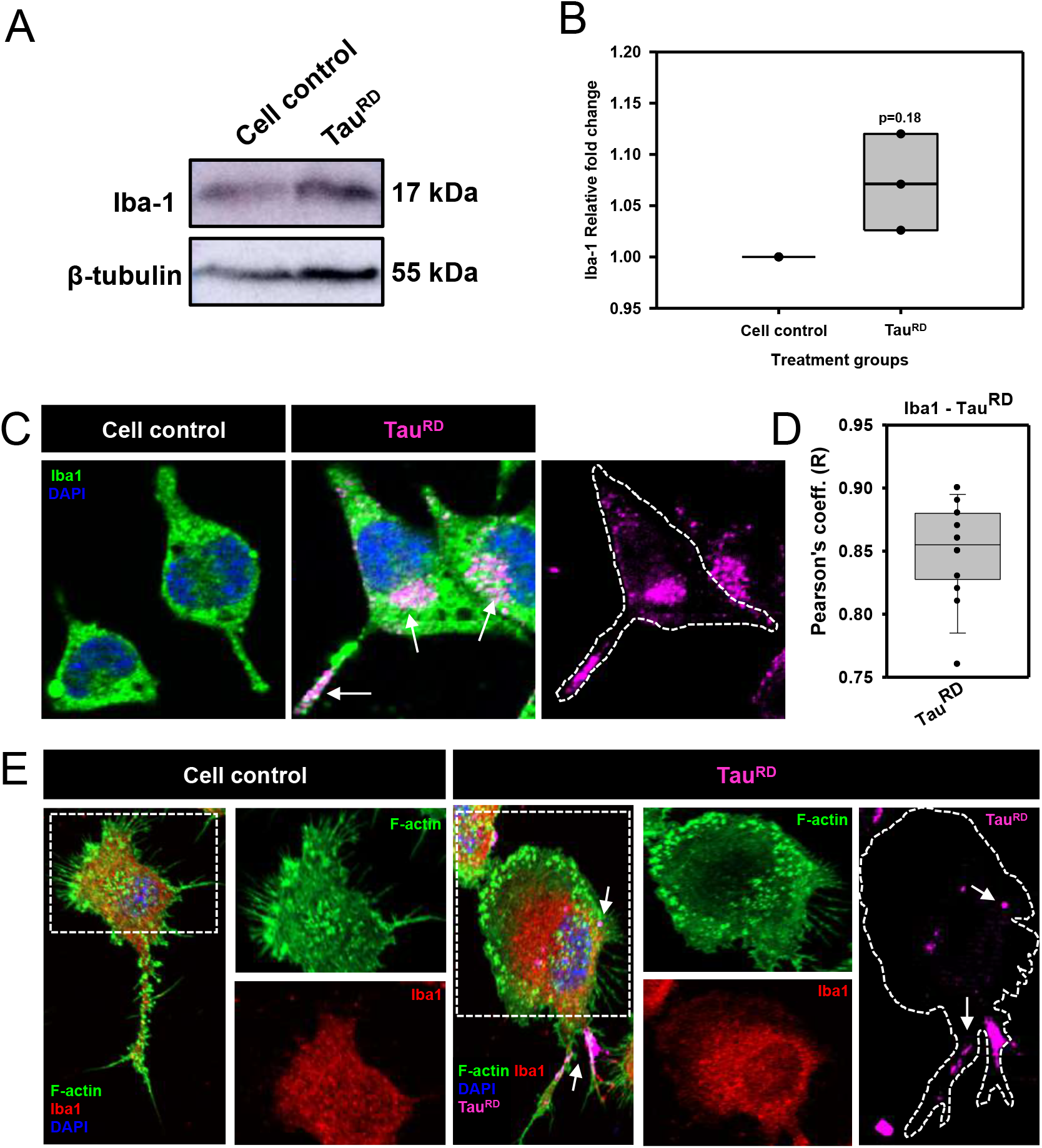
Microglia phagocytosed Tau^RD^ by actin remodeling and colocalized with Iba1. **A.** Upon exposure of Tau^RD^ microglia become activated with an increase in the level of Iba1 by western blot, B. The Iba1 level was quantified from the western blot upon Tau^RD^ exposure as compared to cell control. **C.** Activated microglia internalized Tau^RD^ which were found to be colocalized with Iba1. **D.** Pearson’s coefficient analysis showed that Iba1 were highly colocalized with internalized Tau^RD^. **E.** Activated microglia remodelled membrane-associated actin network with Iba1 colocalization during the Tau^RD^ phagocytosis.

### Phagocytosed Tau^RD^ localize with Iba1 at different cytosolic location

Orthogonal microscopic projections of phagocytic microglia have emphasized that Iba1 was colocalized with internalized Tau^RD^ at different cytosolic location. Phagocytosis of Tau^RD^ was found be mediated from substratum layer through the remodelling of cortical layer actin network. While, the accumulation of Tau^RD^ was found above the plane of cortical layer, at peri-nuclear region (Fig. 6A). The accumulated Tau^RD^ was more colocalized with Iba1 at internal cytosolic layer, but not with membrane-associated actin network (Fig. 6B).

**Figure 6.**
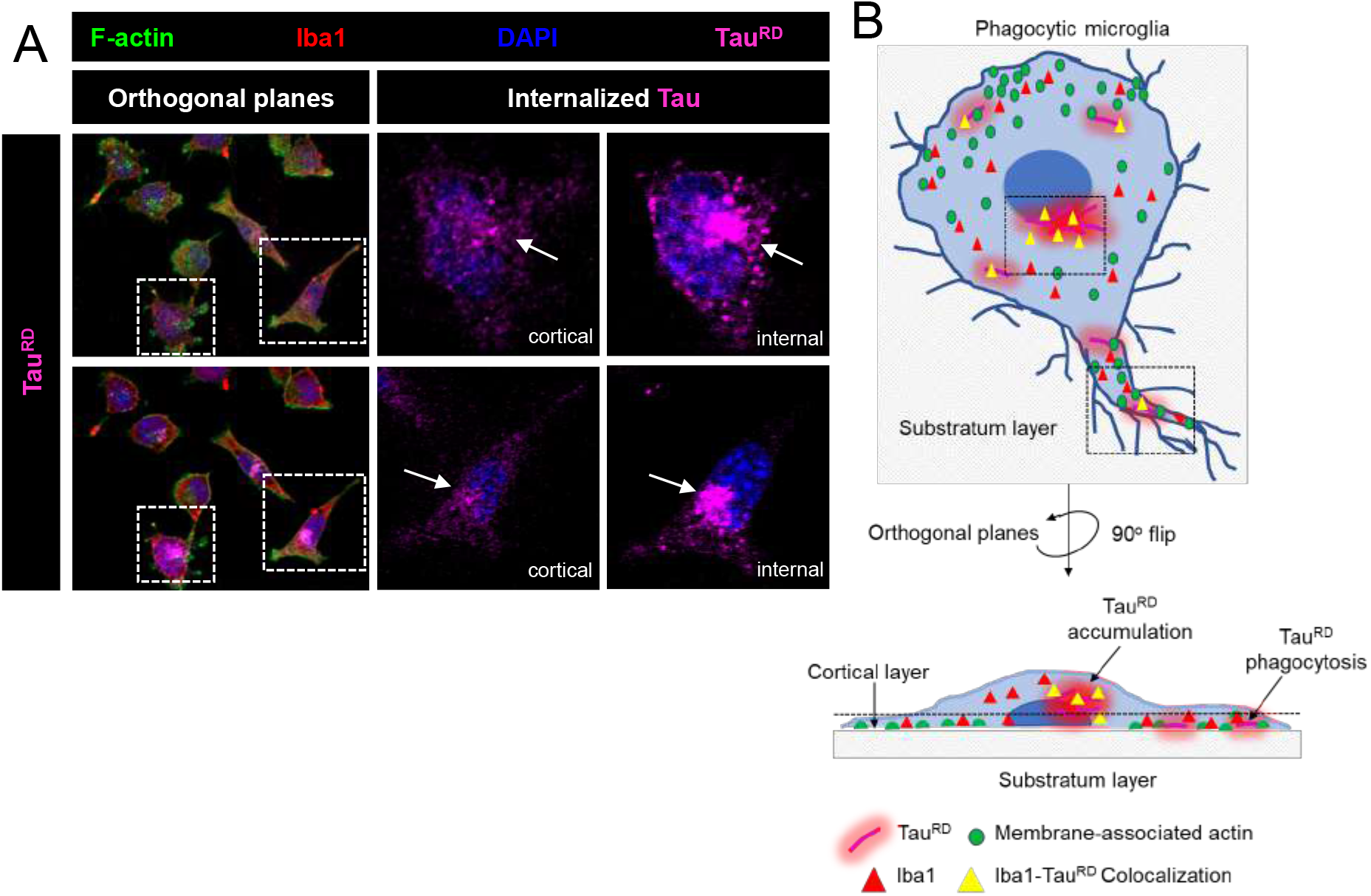
Microglia internalize Tau^RD^ at substratum layer and accumulate at internal cytosolic location. **A.** Phagocytic microglia colocalizes Iba1 with internalized Tau^RD^ at different cytosolic location. **B.** Phagocytosis of Tau^RD^ was occurring from substratum layer through actin remodelling at uropod, while, the Tau^RD^ become accumulated into internal cytosolic location, near to nucleus.

### Microglial P2Y12R interacts with Tau^RD^ and facilitates its internalization

Previously our group showed that full-length Tau interacts with microglial P2Y12R and facilitates its cellular expression. Here, we studied the repeat domain of Tau to emphasize the specific residues involved in P2Y12R interaction. Tau^RD^ was found to be coprecipitated with microglial P2Y12R which ultimately helps in receptor desensitization-mediated Tau internalization (n=3) (Fig. 7A, B). Similarly, when N9 microglia were exposed to Alexa 647-labelled Tau^RD^, P2Y12R along with remodelled actin structures were found to be colocalized with internalized Tau^RD^. While, microglial P2Y12R and Tau^RD^ were colocalized more in cytosolic regions as correlated by positive Pearson’s coefficient (R) values (n=12) (Fig. 7C, D; SI fig. 4A and C). Further, on average 70% of microglia were found to be Tau^RD^ positive, which was ranged from 40 to 100% phagocytosis positive in total population as observed by microscopic fields quantification (n=12) (Fig. 7E).

**Figure 7.**
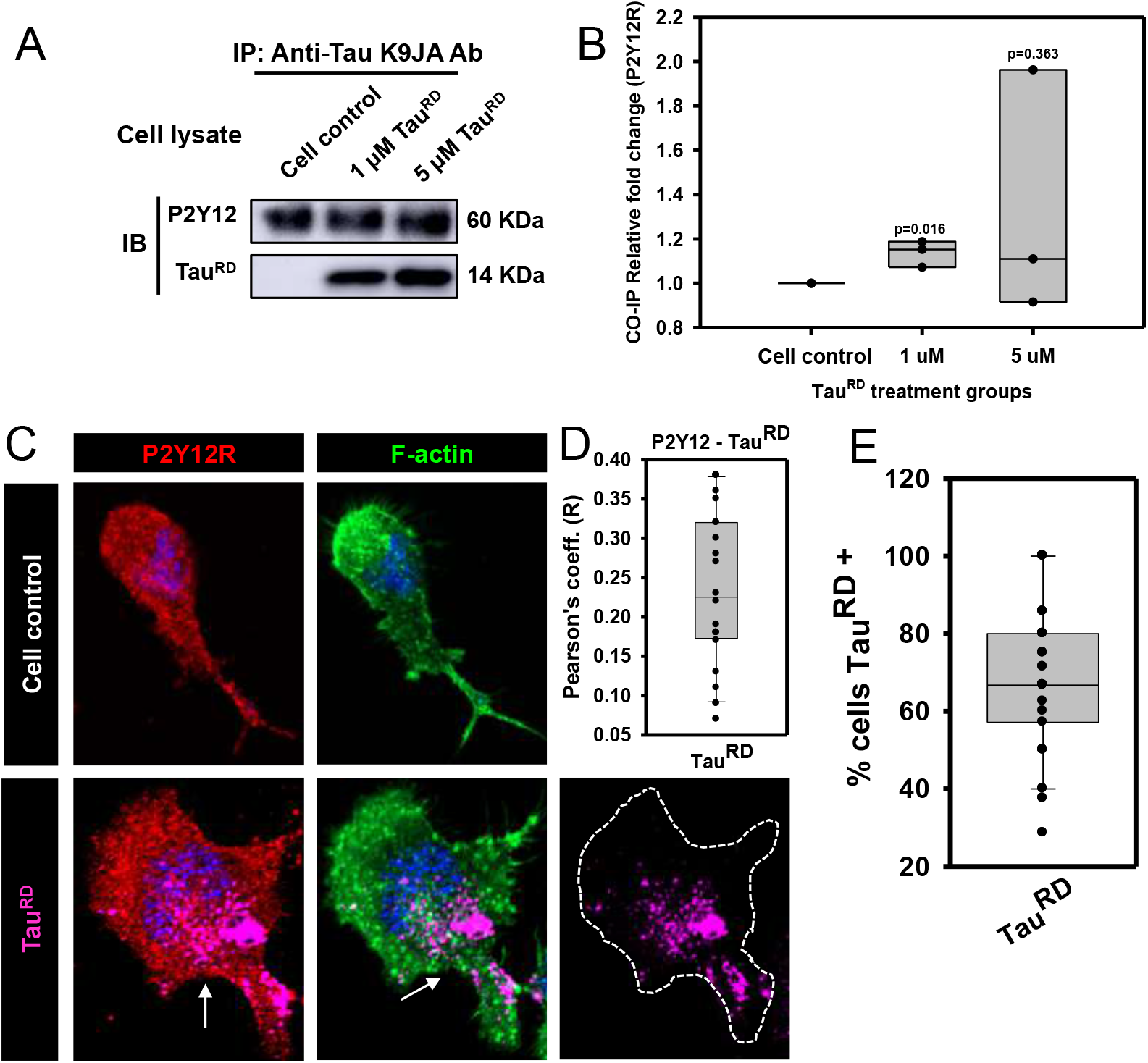
Microglial P2Y12R interacts and internalizes Tau^RD^. **A.** Co-immunoprecipitation performed for extracellular Tau^RD^ and microglial cell lysate from N9 cells against K9JA antibody. Blots were developed against P2Y12 receptor, and K9JA. **B.** 1μM TauRD shows 1.2-fold increase in the band intensity and 5 μM shows 2-folds increase, compared to the cell control. **C, D.** Microglial P2Y12R interacted with Tau^RD^ for internalization which were also colocalized in cytosol which indicate the receptor-mediated phagocytosis. **E.** The % of cells showing phagocytosis of Tau^RD^ were ~70% as found by microscopic images quantification.

### Tau^RD+^ phagocytic microglia remodel P2Y12R-associated filopodia

P2Y12R is a chemotaxis-related purinoceptor which is directly linked with membrane-associated actin-remodelling, adhesion and migration. In our study, we observed that microglia mediated P2Y12R-driven Tau^RD^ phagocytosis where membrane-associated remodelled actin colocalized with Tau^RD^ (Fig. 8A; SI fig. 4B and 5). Similarly, Tau^RD^ exposure has increased the cellular P2Y12R level in microglia by 1.8 folds as compared to cell control (n=3) (Fig. 8B, C). Migratory microglia are known to orchestrate lamellipodia and filopodia during active migration. Moreover, the involvement of podosome is important for matrix adhesion and generation of tensile forces during forward movement. In our study, migratory microglia were observed to form podosome-like structures in frontal lamella and filopodia at uropod during the phagocytosis of Tau^RD^ (Fig. 8D). Also, the filopodia^+^ cells were increased to 70% (some cells 100%) in Tau^RD^ exposed group as compared to untreated control as 40% filopodia^+^ cells (n=12) (Fig 8E). Altogether, resting microglia upon Tau exposure transformed into migratory one with increased accumulation of filopodia along with phagocytic extracellular protein species (Fig. 8F).

**Figure 8.**
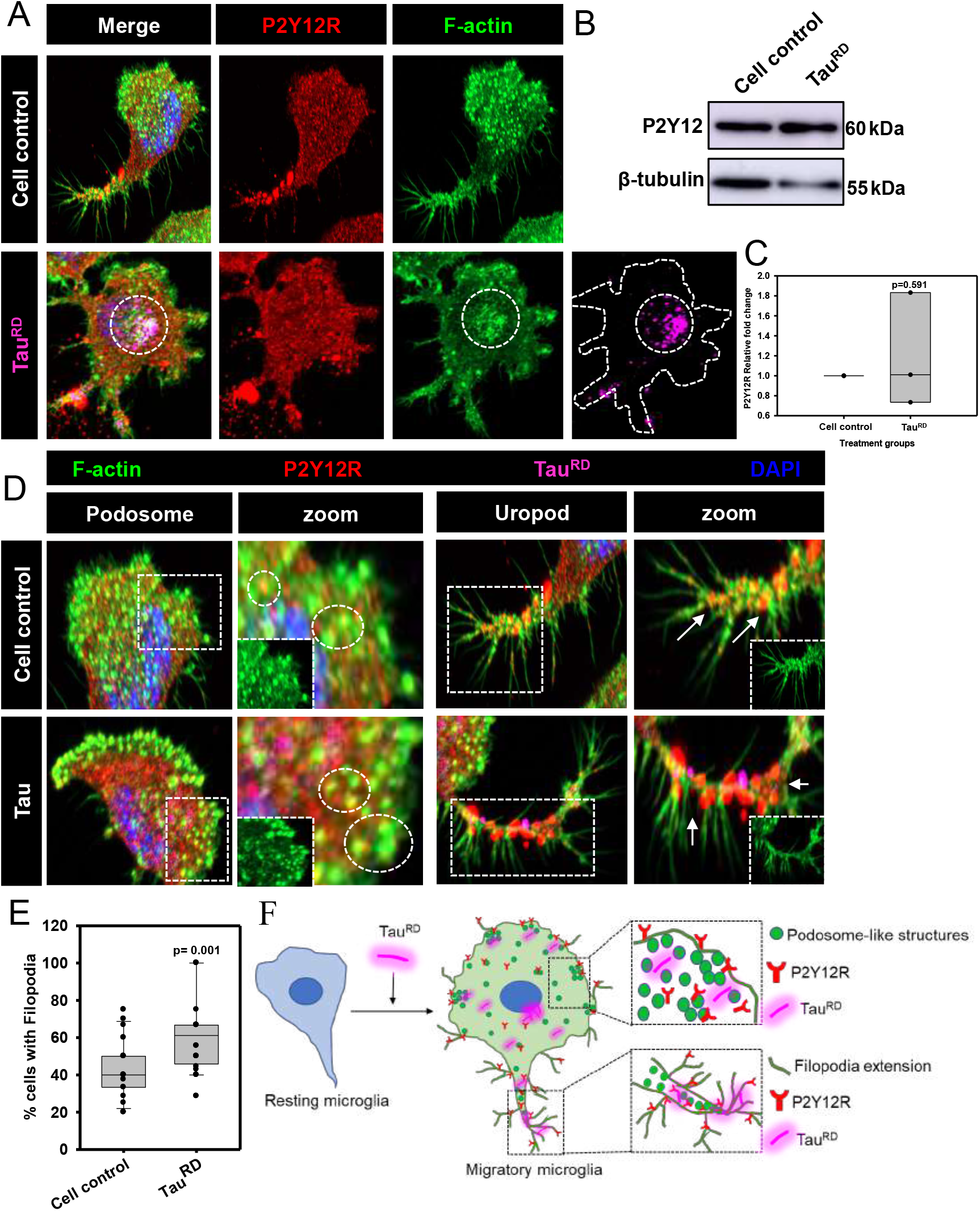
Microglia remodel P2Y12R-localized membrane-associated actin as filopodia by phagocyting Tau^RD^. **A.** Microglia phagocytose Tau^RD^ which were found to be colocalized with podosome-like structures in migratory microglia. **B, C.** Upon extracellular Tau^RD^ exposure, microglial expression level of P2Y12R increased by 1.8-fold than untreated cell control. **D.** Microglia induces the accumulation of P2Y12R-associated filopodia and podosome-like structures in lamellipodia after phagocyting Tau^RD^ as component of active migration. **E.** The % of cells showing filopodia has been increased from 40% to 70% in Tau^RD^ treated group from untreated control. **F.** Tau^RD+^ phagocyting microglia develop branching filopodia and podosome-like actin structures in lamellipodia for P2Y12R-associated migration.

## DISCUSSION

The extracellular Tau and amyloid-β concentration are highly elevated in the AD brain. These intra- and extra-cellular proteins are involved in transmitting intracellular signalling of neuronal and glial cells by interacting with cell-surface membrane receptors. Several GPCRs and other membrane receptors are involved in direct interaction with Tau and amyloid-β proteins (14–17,29). In recent studies, various cell surface receptors are clarified for their role in amyloid interaction especially interaction with extracellular amyloid-β and Tau proteins such as-ApoER, TREM2, scavenging receptors, FcγR, muscarinic acetylcholine receptors, etc (30). In neurons, extracellular Tau has more affinity as compared to acetylcholine towards muscarinic acetylcholine receptors M1 and M3 (16). Hence, extracellular Tau directly interacts with M1 and M3 acetylcholine receptors and increases the levels of intracellular calcium that is toxic to neuronal cells and promotes cell death (16,17,31). In microglia, TREM-2 is a receptor for amyloid-β oligomers, which on its interaction promotes amyloid-β clearance, cytokine synthesis, depolarization, migration, and apoptosis in microglial cells (14). Microglial chemokine receptor, CX3CR1 is recently reported to interact with extracellular Tau and promotes microglial activation and Tau internalization. Tau competes with the CX3CR1 ligand, fractalkine to bind to this receptor and absence of this receptor resulted in impaired microglial activation and Tau internalization (15,20,29). In our previous study, we have reported that the purinergic P2Y12 receptor can directly interact with full-length Tau, promotes microglial activation, mediate actin remodelling for migration and chemotaxis (28). In order to determine that whether Tau^RD^ domain is involved in P2Y12R interaction, we carried out molecular docking and MD-simulation studies with Tau^RD^ and P2Y12R model. Here, we have showed the regionspecific interaction of Tau^RD^ and P2Y12R and its subsequent internalization *via* receptor-mediated endocytosis by various computational, biochemical and cellular studies (Fig. 8). It is also reported that Tau strongly binds to lipid membrane through helices from microtubule-binding repeat regions that leads to membrane disruption and membrane mediated aggregates formation in AD (32,33). Here, we propose that the Tau^RD^-P2Y12R interaction may be supported by cell membrane interactions *i.e*., phospholipids, as it is evident from our analysis (Fig. 2F).

In recent years, the Tau^RD^ domain is attaining more attention in the field of Alzheimer’s disease as it is capable of self-aggregating to form filaments. More-over cross seeding of Tau^RD^ fragments with full-length Tau monomers enhances the rate of aggregation into filamentous structures (34). Aggregated Tau species are capable of propagating in a prion-like fashion among which Tau^RD^ aggregates has been widely studied (35,36). Tau^RD^ transfected HEK cell lines developed pathological features and amyloid conformations of Tau that propagated at least for three generations in mouse models (37,38). Hence the propagation of Tau species from affected to healthy neurons is the main cause of spreading Tauopathy.

But, the glial contribution during Tau propagation needs to be considered thoroughly in due course of disease progression. Recently, it has been found that Tau^RD^ assemblies (n ≥ 3) are the minimum units which are internalized readily by HEK293T cells *via* heparan sulphate proteoglycans (HSPGs) (39). Glial cells *i.e*., microglia and astrocytes do not express Tau, so the presence of Tau in glia signifies only the internalization of extracellular protein and its propagation as seed species (23). In Alzheimer’s disease condition, microglia convert its surveilling state into inflammatory state with active migration, phagocytosis and cytokine production upon encountering extracellular protein deposits (40). During the later stage, excessive extracellular Tau and amyloid-β aggregates may hamper the phagocytic activity of microglia. The dietary fatty acids are playing a significant role in microglial activation and polarization (41). α-linolenic acid treatment is observed to promote the clearance of Tau seeds by enhanced microglial activation, phagocytosis via actin remodelling and degradation of internalized Tau by endosome-lysosome mediated pathway (42,43). Also, a novel antibody targeting Tau^RD^ oligomers can block the intracellular Tau aggregation and also induces cellular uptake-clearance *via* lysosomal pathway in N2A cell model (44). In recent advancements, a neuronal cell model has been established where the internalization of Tau^RD^ and its conversion into endogenous deposits was monitored in real-time. These endogenous deposits are colocalized with P62 and ubiquitin by which Tau^RD^ escapes macroautophagosome pathway and concomitantly propagated to neighbouring neurons and astrocytes *via* contact-mediated tunnelling nanotube (45).

Moreover, in this study, the extracellular Tau^RD^ induces the filopodia extension, colocalized with P2Y12R for dual functioning-directed chemotaxis and receptor-mediated phagocytosis. In activated microglia, phagocytosed Tau^RD^ accumulates at different cytosolic location. In particular, the phagocytosis of Tau^RD^ was evident at uropod, which is associated with branched filopodia. But, the accumulation of Tau^RD^ was occurred at peri-nuclear region with Iba1 colocalization (Fig. 9). Previously, our group emphasized that extracellular Tau are phagocytosed via membrane-associated actin remodelling, such as-lamellipodia and filopodia extension by Iba1^high^ activated microglia (24). Hence, these may signify that membrane-associated actin remodelling especially filopodia formation is necessary for Tau^RD^ engulfment while, the downstream processing may include either degradation or cytosolic accumulation (Fig 9).

**Figure 9.**
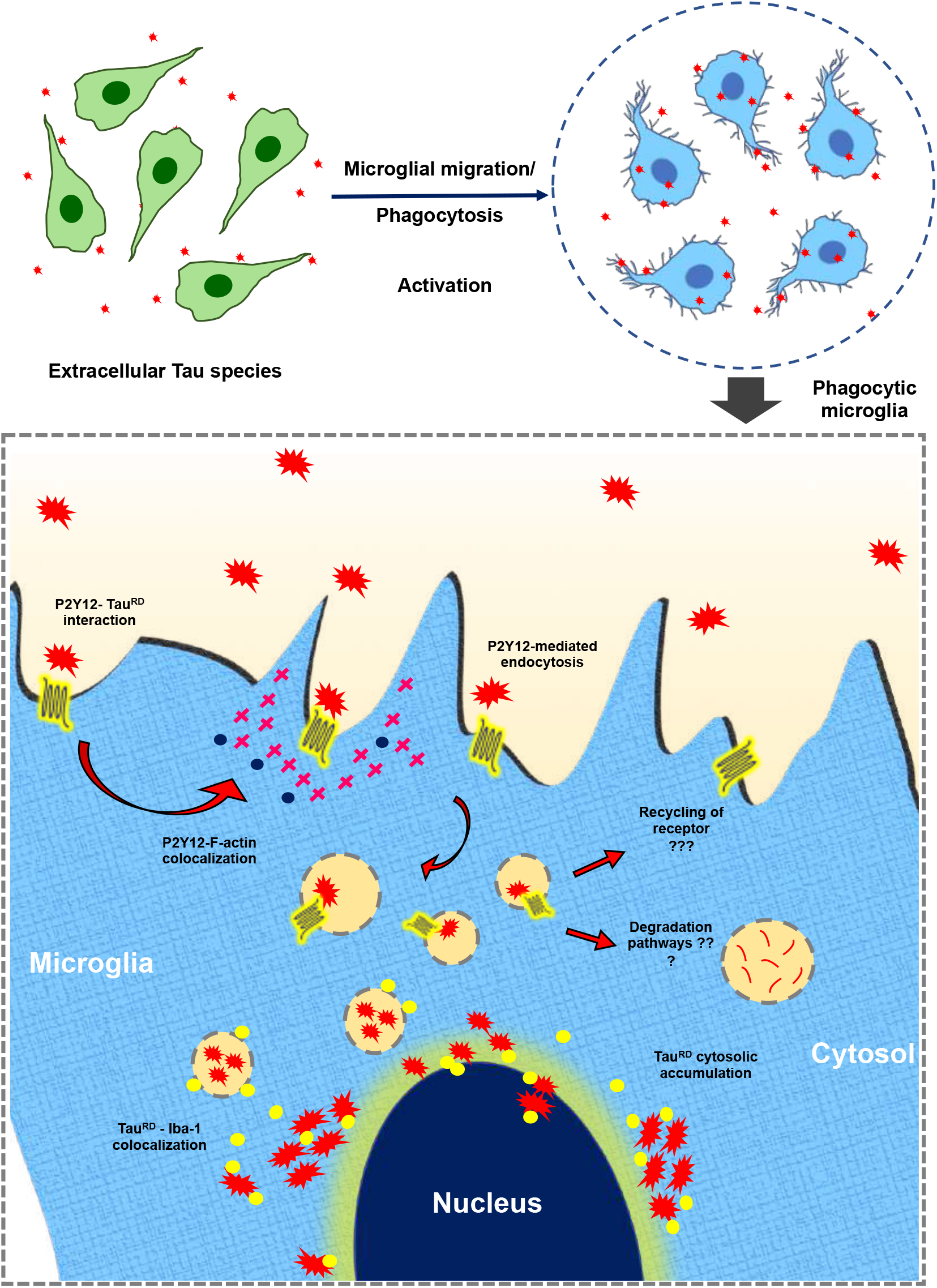
Interaction of Tau^RD^ with microglial P2Y12R for activation and Tau internalization. Extracellular Tau species activate microglia for its active migration and phagocytosis. Tau^RD^ interacts with the extracellular P2Y12 receptor for its internalization mediated by receptor desensitization. The P2Y12 receptor co-localizes with filamentous actin to mediate Tau internalization. The internalized Tau is observed to be co-localized with Iba-1 marker and accumulated in the internal layer near the nuclear region of microglial cells. Hence the microglia is involved in P2Y12R-localized membrane-associated actin remodelling to form filopodial structures in phagocyting Tau^RD^.

## EXPERIMENTAL PROCEDURES

### Chemicals and reagents

For biochemical studies, Luria-Bertani broth from Himedia; ampicillin, KCl, NaCl, Na_2_HPO_4_, KH_2_PO_4_, MgCl_2_, ethylene glycol-bis (b-aminoethyl ether)-N,N,N’,N’-tetraacetic acid (EGTA), phenylmethylsulfonylfluoride (PMSF), ammonium persulfate (APS), ammonium acetate, sodium azide, Dimethyl Sulfoxide Solvent (DMSO) and methanol from MP Biomedicals; isopropyl β-d-1-thiogalactopyranoside (IPTG) and dithiothreitol (DTT) from Calbiochem. CuSO_4_, glutaraldehyde, bicinchoninic acid (BCA), bovine serum albumin (BSA), Thioflavin S (ThS), MES, BES, TritonX-100 and SDS from Sigma; and Tris base, acrylamide, paraformaldehyde, N,N,N’,N’-Tetramethyl ethylenediamine (TEMED), Pierce™ Tris(2-carboxyethyl)phosphine hydrochloride (TCEP-HCl), Pierce™ RIPA buffer, Alexa 647-C2 maleimide purchased from Invitrogen. Immobilon PVDF membrane from Merck. Glycine and Protein Assay Dye Reagent from Bio-Rad. Protease inhibitor cocktail was from Roche cOmplete^TM^. For cell culture, RPMI 1640 media, Fetal Bovine Serum (FBS), trypsin-EDTA, Penicillin-streptomycin, Horse serum were purchased from Invitrogen. In immunofluorescence and western blot studies, we used the following antibodies: total pan-Tau antibody K9JA (Dako, A0024), β-Actin loading control monoclonal antibody (BA3R) (Thermo, MA5-15739), P2Y12R antibody (4H5L19) (Thermo, 702516), Phalloidin-Alexa 488 for F-actin (Thermo, A12379), Goat anti-rabbit IgG (H+L) Cross-adsorbed secondary antibody HRP (Invitrogen, cat no. A16110), anti-mouse secondary antibody conjugated with Alexa flour-488 (Invitrogen, cat no A-11001), Goat anti-rabbit IgG (H+L) Cross-adsorbed secondary antibody with Alexa Fluor 555 (Invitrogen, cat no. A-21428), DAPI (Invitrogen, cat no. D1306), Pierce Co-Immunoprecipitation Kit (Thermo 23600), Precision Plus Protein Standards (Bio-Rad, cat no. 161-0374) and ECL reagent (Bio-Rad, cat no. 1705060). The N9 microglial cell line no. is CVCL_0452.

### Molecular modelling of P2Y12R and repeat domain of Tau

The crystal structures of P2Y12 receptor from RCSB-PDB database (https://www.rcsb.org/) (PDB. ID: 4PXZ and 4NTJ) are used as templates to build the receptor model for active and inactive structures respectively (46–48). SWISS MODEL-server is used to build P2Y12 receptor models (https://swissmodel.expasy.org/) (49). The models with a global model quality estimation (GMQE) of 0.66 is used for further studies. The loops and terminal residues were then built using GalaxyGPCRloop module of Galaxy WEB server (50). The model is then refined using GalaxyRefine module of Galaxy WEB server and the best model is taken for the docking studies (51,52). Model of repeat domain of Tau built by our group in previous studies by Sonawane *et al*. (2019) is adopted for the interaction study (53). For the molecular docking studies, Tau^RD^ is docked to the P2Y12 receptor on the extracellular region that includes the extracellular loops and the N-terminal domain. Molecular docking is performed in ClusPro server where extracellular residues of P2Y12 receptor (referred to UniProt database, Q9H244) are provided for the binding site (54–59). The best model is visualized using PyMol (60) and taken forward for the MD-simulation analysis.

### Molecular Dynamics Simulation of P2Y12R - Tau^RD^ complex

Molecular dynamics simulation of P2Y12 receptor in complex with Tau^RD^ was performed in GROMACS 2019.3 (https://doi.org/10.5281/zenodo.3243833).

Tip3p water model was used for solvation (61–63). GROMOS96 53A6 force field, extended to include Berger lipid parameters was used throughout the simulation (64). Palmitoyl oleoyl phosphatidyl choline (POPC) lipid bilayer model with 128 molecules and their topologies were downloaded from the site of D. Peter Tieleman, Biocomputing group, University of Calgary (http://wcm.ucalgary.ca/tieleman/downloads). The protein-complex was oriented and lipid molecules were packed around the protein using inflateGRO methodology with a scaling factor of 0.95. Energy minimization was performed after every shrinking step with strong position restraints applied to the protein complex. The final membrane density was maintained at ~70 A^2^. The system was then solvated on both hydrophilic sides of lipid bilayer and Cl^-^ ions were added to neutralise the system using genion. The system was then energy minimized using steep descent minimization algorithm. The maximum force was minimized to <1000.0 KJ/mol/nm. Leapfrog integrator was applied to equilibrate the system using NVT and NPT ensembles for 0.1 and 1 ns respectively. Verlet cutoff scheme and PME (particle mesh Ewald) algorithm were used for nonbonded interactions and long-range electrostatics respectively.

Temperature coupling was applied on protein-complex and POPC group and the system was equilibrated to 310 K using V-rescale thermostat (modified Berendsen thermostat). Random velocities were assigned and generated from Maxwell distribution across the periodic boundaries. The pressure is equilibrated at 1.0 bar using Parrinello-Rahman barostat. The production MD was performed for 500 ns (250000000 steps) and observed for the interaction and structural changes in the P2Y12R - Tau^RD^ complex by various methods. RMSD and RMSF graphs were plotted to determine the stability and residual fluctuation during the simulation. The interaction energy between the complex is calculated using g_mmpbsa package of gromacs (65). The 2-dimensional graph of interacting residues is plotted using LigPlot+ (version:2.2) (66). APBS software was used to determine the electrostatic surface potential of the P2Y12R-Tau^RD^ complex as performed by Pontarollo *et al*, 2021 (67). The complex structure obtained after the 500 ns simulation was used for this surface potential analysis. The pqr file was generated for the complex, Tau^RD^ and P2Y12R individually from APBS-PDB2PQR web server using CHARMM forcefield (68). The surface potential was then visualized using PyMol software plugged-in with APBS (69).

### Preparation of Tau^RD^

Tau^RD^ is recombinantly expressed in *E. coli* cells and purified by various chromatographic techniques (70). Tau^RD^ protein is expressed by inducing with 0.5 mM IPTG once the OD (absorbance @ 600 nm) reaches 0.5-0.6. The cells were then harvested following 4 hours of incubation and pelleted down at 4,000 rpm for 10 minutes (4 °C). Resuspended in cell lysis buffer containing 5 mM DTT, 1 mM PMSF and protease inhibitor cocktail. Homogenized at 15,000 psi using cell disruption system. The lysate is then boiled at 90 °C for 15 min by adding 0.5M NaCl and 5 mM DTT. Lysate is cooled and centrifuged at 40,000 rpm for 45 min at 4 °C to remove the precipitated proteins and other cellular components. The supernatant is dialysed overnight with dialysis buffer (added with 50 mM NaCl) and centrifuged again. The supernatant is then loaded for cation exchange chromatography and eluted using buffer containing 1000 mM NaCl. The eluted protein is then loaded for size-exclusion chromatography with 1X PBS buffer in order to get rid of other high and low-molecular weight proteins. Tau^RD^ concentration is then estimated by BCA method.

### Western blot

N9 microglia cells were treated with 1 μM Tau^RD^ and the expression levels of P2Y12R, Iba-1 are determined using western blot analysis. 3 lakh cells were seeded and incubated at 37 °C (5% CO_2_) for 24 hours before the treatment. The treatment was provided for 24 hours and harvested by washing with PBS buffer (pH. 7.4) followed by trypsinization and centrifugation. The cells are lysed with RIPA buffer and the total protein concentration is estimated by Bradford assay. An equal protein volume was used for western blot with rabbit P2Y12R polyclonal antibody (1:1000), Rabbit Iba-1 polyclonal antibody (1:1000) and loading control as mouse β-tubulin monoclonal antibody (1:2000). The quantification of the band intensity was performed using ImageJ 1.53e software. Band intensities are then normalized using house-keeping genes used (β-tubulin) (n=3) and the change in the expression of the protein is compared with the control groups (28).

### Co-immunoprecipitation

N9 microglia cells are seeded in 100 mm petridishes and incubated for 24 hours at 37 °C (5% CO_2_). The cells are then harvested and coimmunoprecipitation was performed using manufacturer protocol with few modifications. Co-IP lysis buffer was used to lyse the cells and incubated for 20 minutes with pipetting at regular intervals. Equal volume of lysate was added with different concentrations of Tau^RD^ (1 μM and 5 μM) and one group without Tau^RD^ as negative control. The mixture is then incubated by constant shaking for 1 hour (RT). Pan-Tau K9JA primary antibody was coupled with the amino-linked resins by reductive amination reaction. Appropriate controls for non-specific binding and isotype IgG were maintained. Tau-lysate mixture is then added to the antibody-coupled resin columns and incubated at 4 °C rotor for overnight binding. The proteins that precipitated along Tau were eluted using the elution buffer according to manufacturer’s protocol. The eluted proteins are analysed by SDS-PAGE and western blot by P2Y12R and K9JA antibody (1:1000 and 1:8000 dilution respectively).

### Alexa Fluor 647 labelling of Tau^RD^ for internalization

100 μM of Tau^RD^ was incubated with 2 μM of Tris(2-carboxyethyl) phosphine (TCEP) in order to maintain a reduced state of sulphur group in Tau cysteine residues. 200 μM of Alexa Fluor™ 647 C2 Maleimide is then slowly added to the protein and incubated for overnight at 4 °C shaker. The untagged molecules are removed by washing with excess PBS buffer using 3 KDa centricons. The final concentration after labelling is measured by BCA assay.

### Immunofluorescence microscopy

The localization of P2Y12R and Iba1 with actin remodelling on microglia upon Alexa647-Tau^RD^ exposure (1 μM) were studied by Immunofluorescence microscopy. N9 cells (10000 cells) were treated with Tau^RD^ monomer for 24 hours. Then, the cells were washed with PBS and fixed with 4% paraformaldehyde solution for 15 minutes and permeabilized with 0.2% TritonX-100. The cells were stained with P2Y12R (1:100), Iba1 (1:100) and phalloidin-alexa488 (1:40) antibody for overnight at 4°C. Then, Alexa flour-secondary antibodies were allowed to bind for 1 hour along with nuclear stain-DAPI (300 nM). The microscopic images were captured in Zeiss Axio observer Apotome2 fluorescence microscope at 63X oil immersion objective. The quantifications were carried out using ZEN 2.3 software. The numbers of filopodia^+^ microglial cells were counted in multiple fields (n=12). The colocalization analysis of Pearson’s coefficient were done by ImageJ software in Tau^RD^ treated group in multiple fields (n=12).

### Statistical analysis

All experiments were performed in two or three biological replicates and triplicate measurements for each individual experiment. One-way ANOVA is used for performing all statistical analyses. The statistical significance of multiple groups has been calculated by Tukey-Kramer’s post-hoc analysis for multiple comparisons at 5% level of significance. The p-values are calculated in comparison with control groups and values are mentioned within the graph. The results are considered as significant if the mean difference between treatment groups is greater than calculated Tukey’s criterion (X-X’>T).

## Data availability

All the data is contained in the manuscript.

## Supporting information

This article contains supporting information.

## Acknowledgements

We thank Shivashankar S and Amit Naglekar for their valuable comments and suggestions in performing the *in-silico* experiments. We are grateful to Chinnathambi’s lab members for their scientific discussions, helpful suggestions and critical reading of the manuscript.

## Author contributions

HC and SC performed the literature search and wrote the manuscript. HC and SC performed the *in-silico* and biochemistry experiments. RD and SC performed the cell biology experiments and wrote the manuscript. SC conceived the idea of the work and supervised the project. All authors read and approved the final manuscript.

## Funding and additional information

This project is supported by in-house CSIR-National Chemical Laboratory grant MLP101726. You can include your fellowships from DBT/UGC.

## Conflict of interest

The authors declare that they have no conflicts of interest with the contents of this article.

## Abbreviations

(AD): Alzheimer’s disease
(APP): amyloid precursor protein
(Aβ): amyloid-β
(APOER): Apolipoprotein E receptor
(CMKLR1): chemokine-like receptor 1
(DAP12): DNAX-activating protein of 12 kDa
(FPR2): formyl peptide receptor 2
(GPCR): G-protein coupled receptor
(GSK-3β): Glycogen synthase kinase −3β
(HEK): human embryonic kidney
(HSPGs): heparan sulphate proteoglycan
(MTOC): microtubule-organizing center
(MAPK): mitogen-activated protein kinase
(POPC): Palmitoyl oleoyl phosphatidyl choline
(ROCK): Rho-associated coiled-coil kinase
(RAGE): receptor for advanced glycation end products
(SCARA1): Scavenger receptor class A member 1
(SCARB1): Scavenger receptor class B member 1
(SYK): spleen tyrosine kinase
(Tau^RD^): Tau-repeat domain
(TLR): Toll-like receptor
(TREM-2): triggering receptor expressed in myeloid cells

